# Evolutionarily Conserved T-tubule-Mitochondria Contacts in Striated Muscle

**DOI:** 10.64898/2025.12.20.694803

**Authors:** Haonan Qi, Le Cao, Ao Li, Qiongqiong Wu, Jing Wang, Mengye Cao, Zhike Liu, Wenjie Sun, Qi Lai, Weijun Pan

## Abstract

The intricate architecture of skeletal muscle is essential for its contractile function, with the spatial organization of organelles such as T-tubules and mitochondria playing a crucial role in maintaining muscle performance and health^1,2^. T-tubules propagate electrical signals deep into muscle fibers to ensure synchronous contraction, while mitochondria provide the necessary energy for muscle contraction^3-5^. Previous studies have extensively documented the interactions between T-tubules and the sarcoplasmic reticulum (SR), which are essential for excitation-contraction coupling^6,7^. However, the detailed spatial relationship between T-tubules and mitochondria has remained relatively unexplored. In this study, we utilized zebrafish and mice as model organisms to investigate the contacts between T-tubules and mitochondria. Using array tomography, deep learning-based analysis, and advanced microscopy techniques, we discovered extensive membrane contact sites, termed T-tubule-Mitochondria (TTM) contacts. These contacts are evolutionarily conserved across vertebrates, including skeletal and cardiac muscles. Our findings reveal a previously unrecognized, intimate spatial coupling between T-tubules and mitochondria, with significant implications for muscle physiology and pathology. Understanding these TTM contacts could provide new insights into the mechanisms underlying muscle function and the development of muscle-related diseases.

## Introduction

The intricate architecture of skeletal muscle is essential for its contractile function, and the spatial organization of its organelles plays a crucial role in maintaining muscle performance and health^1,2,8^. Among these organelles, T-tubules and mitochondria are two key structures. T-tubules, as specialized invaginations of the sarcolemma, are responsible for the rapid propagation of electrical signals deep into the muscle fiber, ensuring synchronous contraction^3,4,9^. Mitochondria, often referred to as the powerhouses of the cell, are critical for energy production, especially in the context of muscle contraction^5,10^. The close spatial relationship between these two organelles is vital for efficient energy transfer and muscle function.

Previous studies have extensively documented the interactions between T-tubules and the sarcoplasmic reticulum (SR), which are essential for excitation-contraction coupling^4,6^. However, the detailed spatial relationship between T-tubules and mitochondria has remained relatively unexplored. Recent advancements in imaging techniques, such as array tomography and focused ion beam-scanning electron microscopy (FIB-SEM), have enabled high-resolution 3D reconstructions of cellular structures^11-13^, providing unprecedented insights into the subcellular architecture of muscle fibers.

In this study, we utilized zebrafish larvae as a model organism to investigate the contacts between T-tubules and mitochondria. Zebrafish larvae offer several advantages, including their small size, which facilitates high-resolution imaging, and their myofibers, which closely resemble the architecture of mammalian skeletal muscle^14,15^. Using a combination of array tomography, deep learning-based analysis, and advanced microscopy techniques, we discovered extensive membrane contact sites between T-tubules and mitochondria, termed T-tubule-Mitochondria (TTM) contacts. These contacts were found to be evolutionarily conserved across vertebrates, including both skeletal and cardiac muscles.

Our findings reveal a previously unrecognized, intimate spatial coupling between T-tubules and mitochondria, with significant implications for muscle physiology and pathology. Understanding these TTM contacts could provide new insights into the mechanisms underlying muscle function and the development of muscle-related diseases.

## Results

### Discovery of T-tubule-mitochondria Contacts

To explore the contacts between mitochondrial reticulum and sarcolemmal system in skeletal muscle, we utilized zebrafish larvae, whose myofibers recapitulate mammalian architecture^15,16^, but with smaller size for vEM-based structural analysis. We generated high-resolution three-dimensional (3D) datasets through array tomography^17,18^, followed by deep learning-based analysis to achieve whole-cell 3D reconstruction of organelle architecture (Fig. 1a, b). We systematically interrogated potential membrane–membrane interactions among 9 reconstructed subcellular compartments within individual myofiber. Beyond confirming previously documented T-tubule–SR junctions^3,19^ and Mito-SR contacts^20-22^ (Fig. 1c), we identified previously unrecognized, extensive membrane contact sites between T-tubules and mitochondria, most residing within contractile units (Fig. 1d). Both volume rendering and axial/transverse orthoslices reveal an intimate, nanometer-scale spatial coupling between T-tubules and mitochondria, which we term T-tubule-Mitochondria (TTM) contacts.

**Figure 1.**
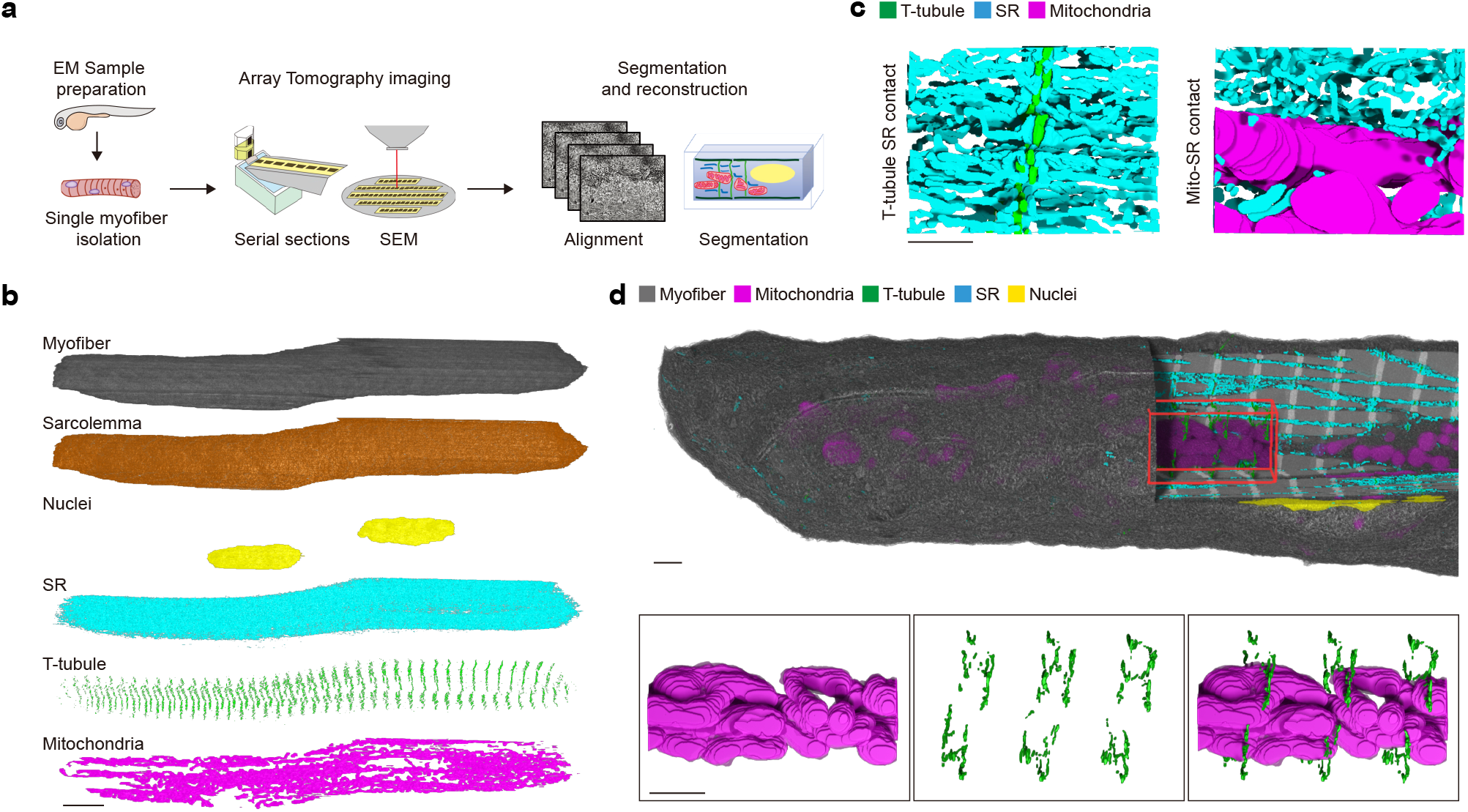
Nanoscale Architecture of T-tubule-mitochondria (TTM) Contacts in Zebrafish Skeletal Muscle. **a**. Schematic of array tomography imaging and analysis approach for zebrafish muscle cell. **b**. 3D reconstruction of organelle classification and segmentation examples from zebrafish skeletal muscle cell. Scale bar, 5 μm. **c**. 3D renderings of T-tubule (green)-SR (blue) and mitochondria (magenta)-SR contacts. Scale bar, 500 nm. **d**. Perspective overview of array tomography volume with segmentation results for mitochondria (magenta), T-tubule (green), nucleus (yellow), and SR (blue) in zebrafish skeletal muscle cell. Illustrative ROI with segmented and 3D rendering of T-tubule and mitochondria. Scale bar, 1 μm.

To verify TTM contacts observed in the vEM, we performed live imaging in zebrafish larvae using muscle-specific *myogenin* promoter-driven expression of EGFP-CAAX (sarcolemma and T-tubule label) and MTS^TOMM20^-mCherry (mitochondrial outer membrane label)^23-25^ (Fig. 2a). Confocal imaging revealed T-tubule with a ladder-like morphology periodically anchored to the intermyofibrillar mitochondria in skeletal muscle (Fig. 2b).

**Figure 2.**
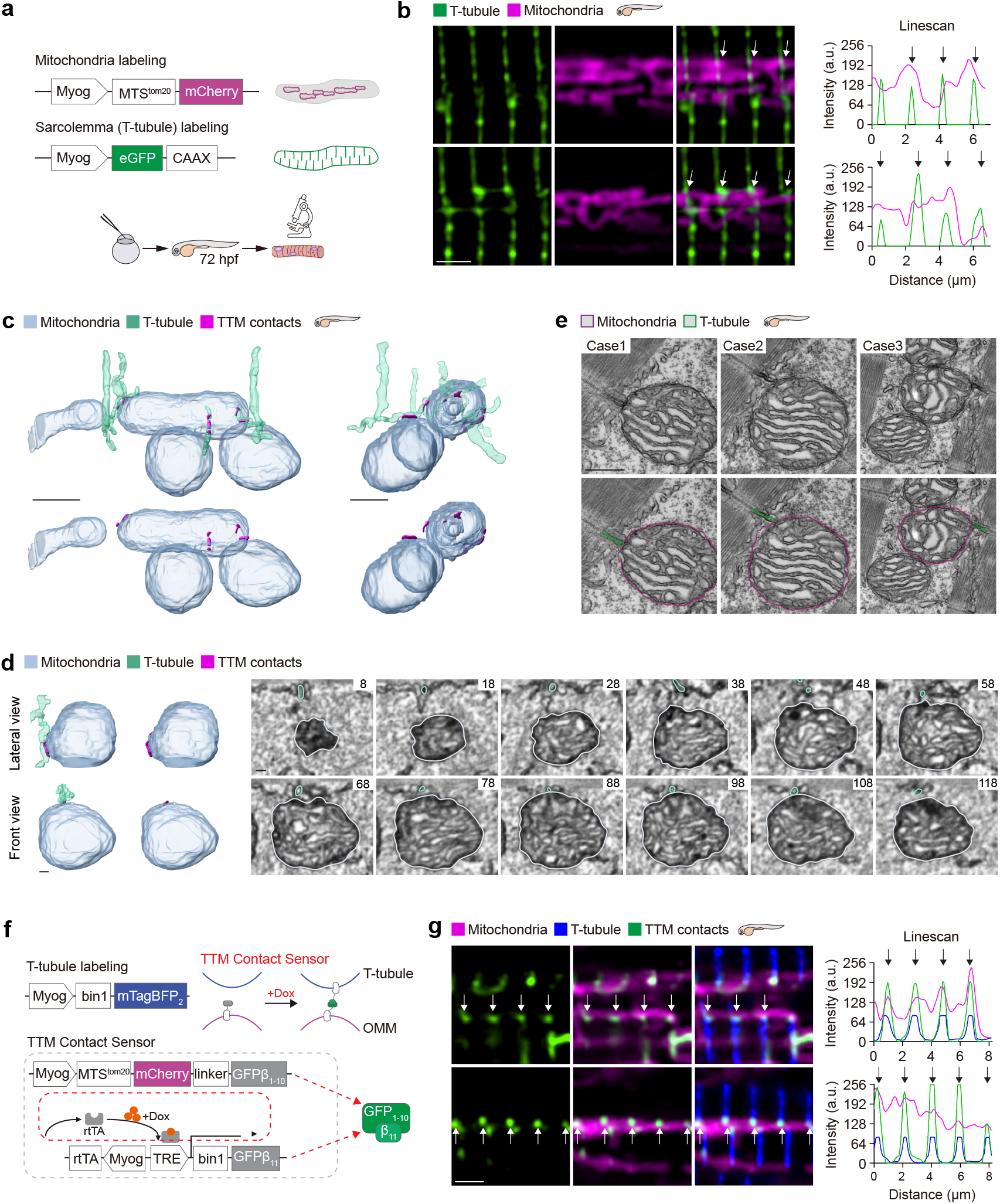
TTM Contacts in Skeletal Muscle. a. Constructs for dual-color imaging to detect T-tubule-mitochondria contacts and workflow for confocal imaging in zebrafish larvae skeletal myofibers, **b**. Confocal image ROI of T-tubule (green) and mitochondria (magenta) in zebrafish larvae skeletal muscle cell. Line scans through mitochondria show relative fluorescence intensity. Scale bar, 2 μm. **c**. FIB-SEM views of T-tubule (green), mitochondria (blue), and TTM contacts (magenta) in zebrafish skeletal muscle cell. Scale bar, 2 μm. **d**. FIB-SEM views of T-tubule (green), mitochondria (blue), and TTM contacts (magenta) in zebrafish skeletal muscle cell. FIB-SEM image series of zebrafish skeletal muscle cell. Scale bar, 100 nm. **e**. TEM images of TTM contacts in zebrafish skeletal muscle cell. Scale bar, 500 nm. **f**. Constructs for T-tubule labeling and inducible TTM contact sensor in a Tet-On system for detecting TTM contact interfaces upon doxycycline treatment, **g** Confocal image ROI of TTM contacts (green), mitochondria (magenta), and T-tubule (blue) in zebrafish skeletal muscle. Line scans show relative fluorescence intensity. Scale bar, 2 μm.

### Ultrastructure and Quantitation of TTM Contacts

To resolve TTM contacts at higher resolution, we leveraged *in situ* analysis on skeletal muscle of zebrafish larvae via focused ion beam-scanning electron microscopy (FIB-SEM), achieving 5 nm isotropic resolution. 3D reconstruction revealed more intimate interactions between T-tubule and mitochondria (Fig. 2c, b). Moreover, transmission electron microscopy (TEM) analysis on isolated single myofiber of zebrafish larvae illustrated T-tubule contacts with mitochondria directly (Fig. 2e).

To further visualize TTM contacts *in vivo*, we targeted split-EGFP fragments^26-28^ to the mitochondrial outer membrane (MTS^TOMM20^) and T-tubules (BIN1)^25,29,30^, respectively. Overexpression artifacts inherent to the split-EGFP-based contact site sensor (SPLICS) were circumvented by integrating the construct into a Tet-On system, yielding temporally precise, low-level expression (Fig. 2f). Live imaging of induced larvae revealed discrete EGFP puncta periodically arrayed along intermyofibrillar mitochondria at T-tubule intersections (Fig. 2g), providing both real-time visualization and quantitative readout of TTM contacts in intact skeletal muscle.

In brief, our discovery of TTM contacts reveal numerous nanoscale bridges that physically couple the T-tubule network to the mitochondrial energy grids, which could have significant implications for muscle physiology and pathology.

### Ultrastructure of Mammalian TTM Contacts

Live-cell imaging of isolated mouse myofibers (CellMask for T-tubule label; PK Mito Deep Red for mitochondria label)^31,32^ and a reanalysis of publicly achieved FIB-SEM datasets^33^ confirmed that TTM contacts are an evolutionarily conserved architectural motif in skeletal muscle across vertebrates (Fig. 3a-c).

**Figure 3.**
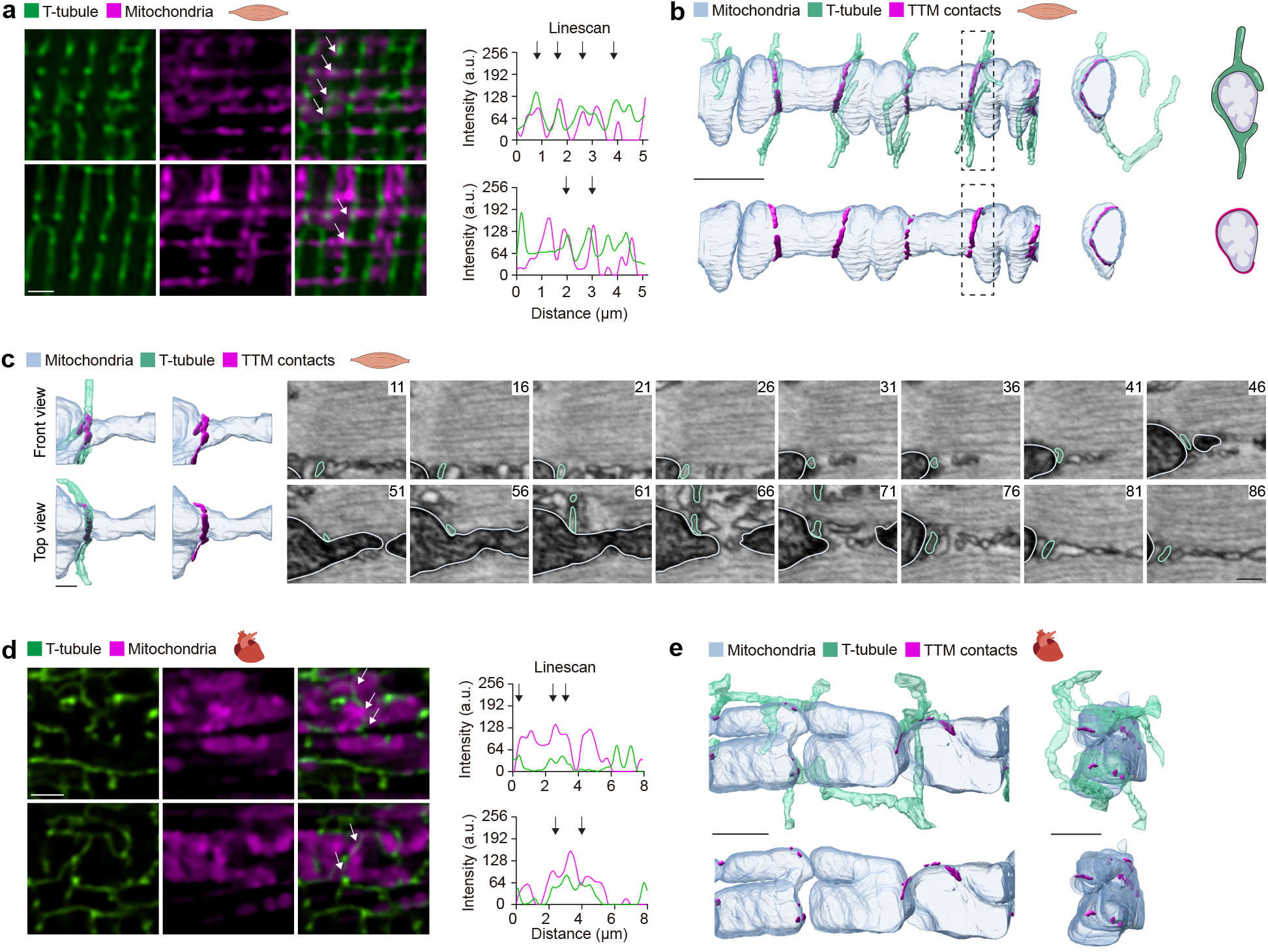
TTM Contacts in Mouse Striated Muscle. **a** Confocal image ROI of T-tubule (green) and mitochondria (magenta) in live mouse soleus muscle fiber. Line scans show relative fluorescence intensity. Scale bar, 1 μm. **b**. FIB-SEM views of T-tubule (green), mitochondria (blue), and TTM contacts (magenta) in mouse skeletal muscle. Scale bar, 1 μm. **c**. FIB-SEM views of T-tubule (green), mitochondria (blue), and TTM contacts (magenta) in mouse skeletal muscle. FIB-SEM image series of mouse skeletal muscle. Scale bar, 200 nm. **d**. Confocal image ROI of T-tubule (green) and mitochondria (magenta) in live mouse cardiomyocytes. Line scans show relative fluorescence intensity. Scale bar, 2 μm. **e**. FIB-SEM reconstruction of T-tubule (green), mitochondria (blue), and TTM contacts (magenta) in mouse cardiac muscle. Scale bar: 1 μm.

Since T-tubule as key component of triad structure in striated muscle^3,34,35^, we performed live-cell imaging of mouse cardiomyocytes (CellMask for T-tubules; PK Mito Deep Red for mitochondria) and reanalysis of public FIB-SEM datasets revealed confirmed similar TTM contacts in cardiac muscle of mammals (Fig. 3d, e). These cardiac observations suggest a fundamental mechanism conserved across striated muscles.

To systematically chart the spatial distribution of TTM contacts and determine their precise alignment with the skeletal-muscle mitochondrial reticulum, we performed a network-scale interrogation of isotropic 10-nm FIB-SEM volumes that encompass the entire intermyofibrillar mitochondria (Fig. 4a). Intermyofibrillar mitochondria adopt three stereotyped topologies. Parallel mitochondria (P-Mito) run longitudinally along the contractile axis, spanning successive I- and A-bands as continuous, fused tubules. Vertical mitochondria (V-Mito) are transversely oriented, anchored orthogonally at the Z-lines. Connective mitochondria (C-Mito) form slender bridges that knit P-Mito and V-Mito into a cohesive reticulum (Fig. 4b-f). TTM contacts are overwhelmingly concentrated on P-Mito, generating a quasi-crystalline array with a periodic spacing of ∼0.9 µm along each P-Mito segment (Fig. 4g-i). Remarkably, virtually every TTM contact lies adjacent to the filament sliding zone (Fig. 4j) and refined measurements reveal that the mean distance from each TTM contact to the nearest contractile unit is less than 20 nm (Fig. 4k) placing them immediately next to the ATP hydrolysis site by myosin motors^36^.

**Figure 4.**
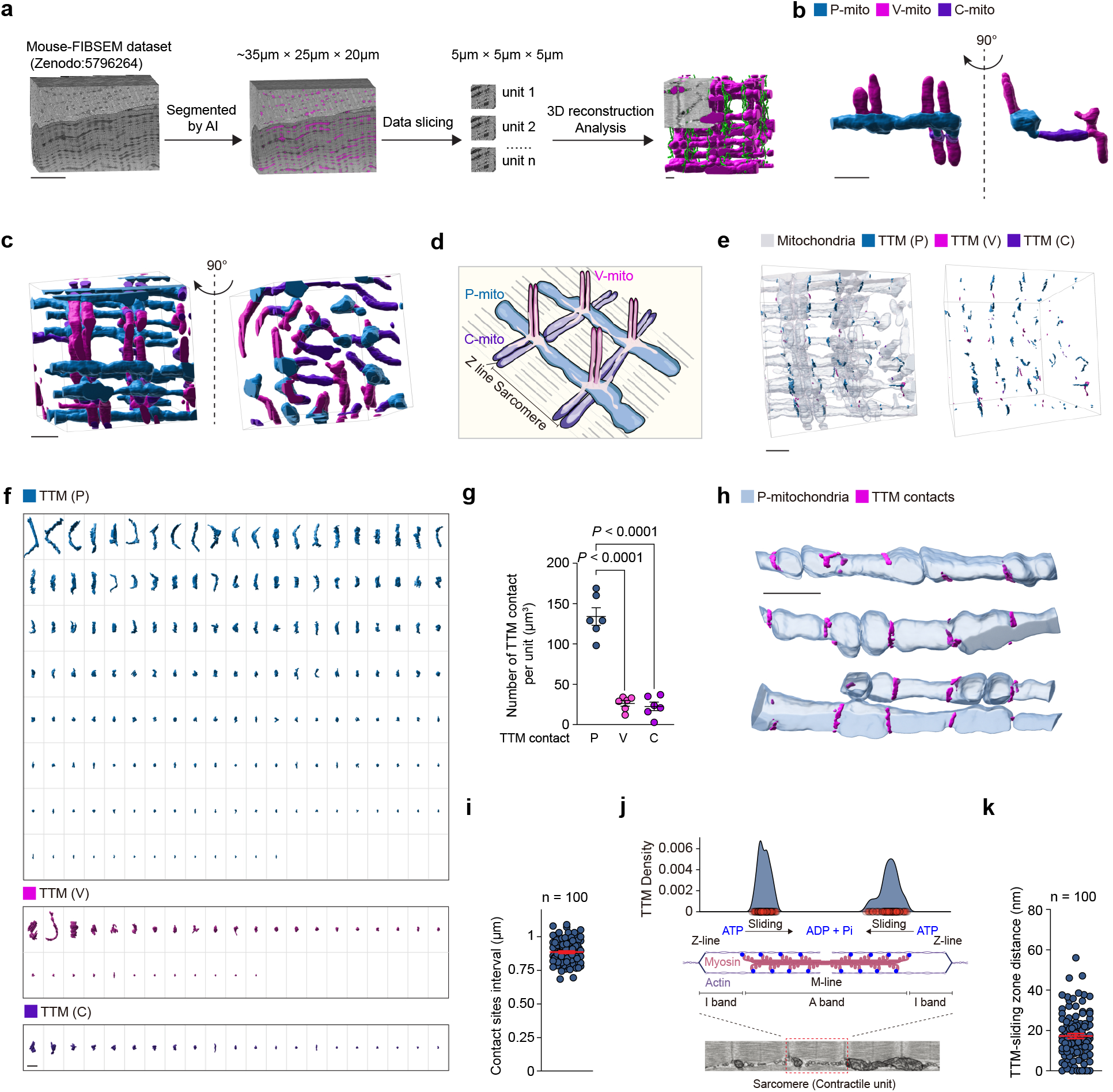
TTM Contacts Locate in Proximity to the Contractile Machinery. **a** Workflow for segmentation and classification of mitochondrial networks in mouse skeletal muscle FIB-SEM data. **b**. 3D views of P-Mito (blue), V-Mito (magenta), and C-Mito (purple) in mouse skeletal muscle. Orthogonal view shown. Scale bar, 1 μ m. **c**. Extended 3D views of P-Mito, V-Mito, and C-Mito in mouse skeletal muscle. Orthogonal view shown. Scale bar, 1 μ m. **d**. Schematic depiction of P-Mito, V-Mito, and C-Mito based on orientation relative to mouse skeletal muscle fibers. **e** 3D views of mitochondria (grey) and TTM contacts across mitochondrial subtypes in mouse skeletal muscle. Subtype-specific TTM contacts shown, **f**. Classification of TTM contacts in a gallery model, **g** Quantitative analysis of TTM contact number in P, V and C-Mito within mouse skeletal muscle FIB-SEM imaging, n = 6 cases. **h** Longitudinal views of periodic TTM contacts along P-Mito in mouse skeletal muscle FIB-SEM datasets. Scale bar, 1 μ m. **i**. Average TTM contact sites interval along P-Mito (0.9 ± 0.1 μ m). Data are mean ± s.e.m., n = 100. **j**. Density distribution of TTM contacts in mouse skeletal muscle sarcomere, n = 75 data point, **k**. Average distance between TTM contact and the sliding zone. Data are mean ± s.e.m., n = 100.

Thus, these TTM contacts are, on one hand, part of the terminal branches of the periodically arranged T-tubule system derived from the sarcolemma, and on the other hand, bridge energy production (mitochondria) and energy utilization (filament sliding zone) in the sarcomeres.

## Discussion

The discovery of T-tubule-Mitochondria (TTM) contacts in skeletal muscle represents a significant advancement in our understanding of the subcellular architecture and functional organization of muscle fibers. Our findings reveal an extensive and intimate spatial coupling between T-tubules and mitochondria, which has important implications for muscle physiology and pathology.

The presence of TTM contacts in both zebrafish and mammalian skeletal muscle, as well as in cardiac muscle, highlights the evolutionary conservation of this structural motif. This conservation suggests that TTM contacts play a fundamental role in the function of striated muscles across vertebrates. The proximity of T-tubules to mitochondria, particularly within the contractile units, indicates that these contacts may facilitate efficient energy transfer from mitochondria to the sites of ATP utilization during muscle contraction. This spatial arrangement could optimize the delivery of ATP to myosin motors, thereby enhancing muscle performance.

The detailed ultrastructure of TTM contacts, as revealed by our high-resolution imaging techniques, provides new insights into the spatial organization of these interactions. The periodic arrangement of TTM contacts along P-Mito and their proximity to the filament sliding zone suggest a highly organized and functional architecture. This quasi-crystalline array of TTM contacts, with a periodic spacing of approximately 0.9 µm, may ensure that energy production and utilization are spatially coordinated, minimizing energy loss and maximizing contractile efficiency.

The use of advanced imaging techniques, such as array tomography, focused ion FIB-SEM, and TEM, has been instrumental in uncovering these previously unrecognized contacts. The integration of deep learning-based analysis for whole-cell 3D reconstruction further enhances our ability to visualize and quantify these interactions. Additionally, the application of SPLICS in live imaging experiments provides real-time visualization and quantitative analysis of TTM contacts in intact muscle tissue. These methodological advances open new avenues for studying organelle interactions in muscle and other cell types.

The identification of TTM contacts may have significant implications for understanding muscle-related diseases. Disruptions in the spatial organization of T-tubules and mitochondria have been implicated in various muscle pathologies, including muscular dystrophies and cardiomyopathies. Further investigation into the role of TTM contacts in these conditions could provide new therapeutic targets and diagnostic markers. For instance, interventions aimed at restoring or enhancing TTM contacts might improve muscle function in patients with these disorders.

Future research should focus on elucidating the molecular mechanisms underlying TTM contacts. Identifying the proteins and signaling pathways involved in the formation and maintenance of these contacts could provide a deeper understanding of their function. Additionally, studies examining the role of TTM contacts in different physiological and pathological conditions, such as aging, exercise, and muscle injury, could reveal their dynamic nature and potential for therapeutic modulation.

In conclusion, our discovery of TTM contacts in skeletal and cardiac muscles highlights a previously unrecognized aspect of muscle cell biology. These contacts represent a critical structural and functional link between energy production and utilization in muscle fibers, with broad implications for muscle physiology and pathology. Further exploration of TTM contacts will undoubtedly contribute to our understanding of muscle function and the development of novel therapeutic strategies for muscle-related diseases.

## Materials and Methods

### Animal Husbandry

Zebrafish were maintained according to standard protocols and handled in compliance with the guidelines of the Institutional Animal Care and Use Committee at the Shanghai Institutes for Nutrition and Health, Chinese Academy of Sciences. Larvae were incubated at 28.5°C and treated with 0.003% 1-phenyl-2-thiourea (Merck) from 12 hours post fertilization (hpf) to inhibit pigment formation. The Tubingen wild-type (WT) strain was used in this study.

Mice were housed under specific pathogen-free conditions and handled in accordance with the guidelines of the Institutional Animal Care and Use Committee at the Shanghai Institutes for Nutrition and Health, Chinese Academy of Sciences (SINH-2025-PWJ-1).

### Plasmid Construction and Microinjection

Split-GFP-based contact site sensor (SPLICS) plasmids were constructed as follows: pTol2-myogenin:rtTA; TRE:bin1-GFP_b11_; pTol2-myogenin:MTS^tomm20^-mCherry-GFP_b1-10_. These constructs were optimized using the Tet-ON system to induce lower-dose expression. Other constructs included: pTol2-myogenin:bin1-mTagBFP2; pTol2-myogenin:eGFP-caax; pTol2-myogenin:MTS^tomm20^-mCherry. All constructs were confirmed by DNA sequencing.

For zebrafish transient transgenesis, purified plasmids (15 pg) were injected into one-cell-stage larvae with tol2 transposase mRNA (15 pg).

### Isolation of Zebrafish Larvae Skeletal Myofibers

Zebrafish larvae at 3 dpf were anesthetized in tricaine (Merck). The heads of the larvae were removed with a scalpel, and the remaining tissue, primarily skeletal muscle, was transferred into a 1.5 ml centrifuge tube. The tissue was washed twice with sterile phosphate-buffered saline (PBS, Gbico), and 600 μl of 0.25% trypsin (Gbico) was added to the tube containing the sample. The sample was digested for 30 minutes at 28.5°C, with trituration every 10 minutes using a P1000 pipette (Gilson). The tubes were then centrifuged at 700-800 rpm for 5 minutes to pellet the cells. The supernatant was removed, and the cells were washed twice with CO_2_-independent media (Gbico). The cells were resuspended in 600 μl of fresh CO_2_-independent media and passed through a 70 μm filter (Miltenyi Biotec) to remove debris. The myofiber suspension was then added to poly-L-lysine-coated coverslips.

### Isolation of Adult Mice Single Muscle Fiber

Intact soleus muscles were isolated from tendon to tendon from 8-week-old mice. The excised muscles were incubated in dissociation medium (DM) containing DMEM (Invitrogen), gentamycin (50 µg ml^-1^, MCE), FBS (2%, HyClone), and collagenase A (4 mg ml^-1^, Roche) at 37°C in a CO_2_ incubator for 2 hours. The tissues were then transferred to incubation medium (IM) composed of DMEM, gentamycin (50 µg ml^-1^), and 10% FBS. Single myofibers were obtained by gently flushing the muscle with a large-bore pipette. Long, non-contracted fibers were selected for further analysis.

Single cardiomyocytes were isolated using a Langendorff apparatus^37^. Briefly, the heart from an 8-week-old mouse was rapidly excised, and the proximal aorta was cannulated with a 21-gauge needle connected to a Langendorff perfusion system. The heart was perfused at a constant flow rate of ∼4 ml min^-1^ with Tyrode solution for 4 minutes, followed by enzymatic digestion with Collagenase buffer containing Collagenase Type II (1 mg ml^-1^, Worthington), Protease Type XIV (0.1 mg ml^-1^, Merck), and 0.05 mM CaCl_2_ in Tyrode solution for ∼30 minutes. Following digestion, cardiomyocytes were dissociated by gentle pipetting, and enzymatic activity was quenched by adding 3 ml of stopping buffer containing 10 mg ml^-1^ BSA in Tyrode solution. The isolated cardiomyocytes were sequentially washed and sedimented twice with Ca^2+^ solution I (200 µM CaCl_2_ in stopping buffer) and Ca^2+^ solution II (500 µM CaCl_2_ in stopping buffer). Cells were then resuspended in 10 ml of plating medium composed of M199 (Gibco) supplemented with 5% FBS and 10 mmol L^-1^ BDM (Merck), seeded onto Laminin-coated Petri dishes (Cellvis), and incubated at 37°C in a CO_2_ incubator for 2 hours prior to imaging.

### Light Microscopy and Analysis (2 Parts)

#### Part 1 Confocal Microscopy

Confocal imaging was performed using an IXplore SpinSR microscope (Olympus) equipped with a 60× silicone oil-immersion objective or a 100× oil-immersion objective, with consistent imaging parameters. The 405 nm laser was utilized to detect bin1 (T-tubule)-mTagBFP2. The 561 nm laser was employed to detect MTS^tomm20^ (OMM)-mCherry. The 488 nm laser was used to detect splitGFP (contact sensor) signals and EGFP-CAAX (sarcolemma), while the 647 nm laser was used to detect Alexa 647. Confocal images were processed using Fiji and Imaris (Oxford Instruments) software.

#### Part 2 Mice Single Muscle Fiber Staining and Imaging

For mitochondria and T-tubule labeling in single skeletal and cardiac muscle fibers, isolated muscle fibers were incubated in incubation medium (IM) supplemented with CellMask™ Green (1×, Invitrogen) and PK Mito Deep Red (250 nmol ml^-1^, Genvivo) for 15 minutes to label T-tubules and mitochondria, respectively. Following staining, fibers were washed three times with IM and transferred to a Petri dish for confocal imaging. Imaging was performed using a IXplore SpinSR microscope (Olympus) equipped with a 100×, 1.5 NA oil-immersion objective. A 488 nm laser was used to detect CellMask™ Green, while a 640 nm laser was used to excite PK Mito Deep Red. Image processing and rendering were conducted in Imaris (Oxford Instruments), and lipid droplet number and diameter were analyzed using Fiji.

### Electron Microscopy and Analysis (5 Parts)

#### Part 1 Sample Preparation

Zebrafish skeletal myofibers were dissociated and settled for approximately 1 hour onto poly-L-lysine-coated plastic coverslips at room temperature. The coverslips with myofibers were fixed in a fixative solution (2.5% glutaraldehyde in 0.1 M phosphate buffer, pH 7.2) for 2 hours at room temperature. The myofibers were then transferred to fresh fixative solution and fixed overnight at 4°C. Samples were washed six times with 0.1 M phosphate buffer for 5 minutes each at 4°C. Subsequently, samples were post-fixed in 1.5% potassium ferrocyanide (K_3_[Fe(CN)_6_]) in 0.1 M phosphate buffer with 1% osmium solution (OsO_4_) for 10 minutes at 4°C, washed six times in bi-distilled H_2_O for 5 minutes each, and incubated in fresh thiocarbohydrazide (TCH) solution (1% TCH in bi-distilled H_2_O) for 30 minutes at 37°C. After six washes with bi-distilled H_2_O at room temperature, samples were secondarily post-fixed in 1% osmium solution for 30 minutes at 4°C and washed six times in bi-distilled H2O. Next, the samples were incubated in 2% uranyl acetate solution at 4°C overnight, washed six times in bi-distilled H_2_O at room temperature. The samples were then dehydrated sequentially with gradient-diluted ethanol solutions (30%, 50%, 70%, 80%, 90%, 100%, and twice 100%; 5 minutes each) at room temperature. The samples were treated with propylene oxide three times for 5 minutes each, incubated in freshly made 50% Epon (50% propylene oxide) for 2-3 hours, replaced with 75% Epon for 2-3 hours at room temperature, and finally incubated with fresh 100% Epon overnight at room temperature. The next day, samples were transferred to fresh 100% Epon for another 1-hour treatment. Excess resin was removed, and the samples were transferred to a 60°C oven for polymerization. Part of the samples were placed on aluminum ZEISS SEM mounts and polymerized at 60°C for 2 days, then gold-coated for FIB-SEM imaging. For other samples, after dividing into small pieces, they were resin-embedded and polymerized at 60°C for 2 days, and then resin-embedded blocks were subjected to ultramicro serial sectioning for Array Tomography imaging and TEM imaging.

#### Part 2 TEM Imaging

Samples were prepared as described above. After incubating at 60°C for 2 days, resin-embedded blocks were sectioned in an ultramicrotome (Leica) to obtain 50 nm sections. Samples were stained with uranyl acetate/lead citrate, and high-resolution images were acquired using a JEOL-1230 80kV electron microscope (JEOL) and a Talos L120C electron microscope (Thermo Fisher Scientific).

#### Part 3 Array Tomography Imaging

Samples were prepared as described above. After incubating at 60°C for 2 days, resin-embedded blocks were sectioned in an ultramicrotome (Leica) to obtain 70 nm sections, and serial sections were manually collected on silicon wafers. Array Tomography images were acquired using a ZEISS Gemini 300 with ZEISS Atlas 5 software (Carl Zeiss Microscopy GmbH, Jena, Germany). For analysis, one complete muscle cell was imaged in 180 serial sections, with a field of view of 15,000 × 5,000 pixels on each section at 5 nm pixel^-1^.

#### Part 4 FIB-SEM Imaging

FIB-SEM images were acquired using a Thermofisher Helios CX with Auto slice and view software and collected using an in-column BSE detector. Images were acquired at 2 keV, 0.2 nA current at a working distance of 2.5 mm. Images were acquired at 5 nm pixel-1 with a 4 µs dwell time. FIB milling was performed at 30 keV, 2.5 nA current, and 5 or 10 nm thickness. Image stacks within a volume were exported as TIFF files using Auto slice and view software for analysis.

#### Part 5 Image Segmentation and 3D Reconstruction

Prior to segmentation, alignment of Array Tomography volumes and FIB-SEM volumes was performed using the Registration plugin and the TrakEM2 plugin in Fiji. Image segmentation of organelle structures was conducted using the Segmentation Wizard module and Deep Model Training in Dragonfly (2022.2) software. For analyses involving different organelle structures, training classes were created for mitochondria, sarcolemma, T-tubule, lipid droplets, sarcoplasmic reticulum, sarcomeric I-bands, A-bands, and nucleus when visible. After initial training, the results of the initial segmentation were displayed for checking for mislabeling errors, and additional training iterations were performed when necessary to minimize errors. 3D reconstruction and animations of the skeletal muscle cell, organelle structures, and T-tubule-mitochondria contacts were generated in Dragonfly (2022.2) and rendered in Imaris (Oxford Instruments).

### Statistics and Reproducibility

Statistical analyses were performed using GraphPad Prism (version 10) or R (version 4.4.1). Comparisons between two groups were assessed using two-tailed Student’s t-tests, while one-way ANOVA was applied for multiple group comparisons, as appropriate. Data are presented as mean ± s.e.m., and the number of biological replicates (n) is specified in the corresponding figure legends. All experiments were independently repeated at least twice to ensure reproducibility. No statistical methods were used to predetermine sample size.

## Acknowledgements

We are grateful to X.W. (Electron Microscopy Core Facilities of ION, CAS), Y.S. (Zeiss) and R.M. (Thermo Fisher Scientific) for assistance with array tomography and FIB-SEM imaging in zebrafish. We thank J.W. (Fudan University) for technical support with myofiber isolation, and Q.L. from the H.Y. laboratory (SINH, CAS) for assistance with cardiomyocyte isolation. We also thank Z.W., J.Z., T.Z. and Z.G. for the constructive suggestions during our manuscript preparation. We acknowledge the use of the animal facilities at the Shanghai Institute of Nutrition and Health, Chinese Academy of Sciences.

The work was supported by a grant from National Key R&D Program of China (2023YFA1802000), Shanghai Pilot Program for Basic Research-CAS Shanghai Branch (JCYJ-SHFY-2022-006), SINH Research Project (JBGSRWBD-SINH-2021-04) and National Natural Science Foundation of China (32130033 and 31571505) to W.P.

## Author contributions

H.Q., L.C. and W.P. developed the concepts and designed the experiments. H.Q., L.C. planned and performed all the experiments and analyzed data. A.L. and Q.W. assisted with plasmid construction. J.W. assisted with intramuscular AAV injection. A.L., M.C., Z.L., W.S. and Q.L. assisted with mitochondrial segmentation of mouse skeletal muscle FIB-SEM datasets. H.Q. and L.C. made figures and models. H.Q., L.C., and W.P. wrote the manuscript. W.P. supervised the project.

## Data and Code availability

All data supporting the results of this study are available from the corresponding authors upon reasonable request. There is no original code generated in this study.

## Reference

1. Frontera, W.R., and Ochala, J. (2015). Skeletal muscle: a brief review of structure and function. Calcif Tissue Int 96, 183–195. 10.1007/s00223-014-9915-y.

2. Glancy, B., and Balaban, R.S. (2021). Energy Metabolism Design of the Striated Muscle Cell. Physiol Rev. 10.1152/physrev.00040.2020.

3. Al-Qusairi, L., and Laporte, J. (2011). T-tubule biogenesis and triad formation in skeletal muscle and implication in human diseases. Skelet Muscle 1, 26. 10.1186/2044-5040-1-26.

4. Kuo, I.Y., and Ehrlich, B.E. (2015). Signaling in muscle contraction. Cold Spring Harb Perspect Biol 7, a006023. 10.1101/cshperspect.a006023.

5. Glancy, B., Hartnell, L.M., Malide, D., Yu, Z.X., Combs, C.A., Connelly, P.S., Subramaniam, S., and Balaban, R.S. (2015). Mitochondrial reticulum for cellular energy distribution in muscle. Nature 523, 617–620. 10.1038/nature14614.

6. Xu, J., Liao, C., Yin, C.C., Li, G., Zhu, Y., and Sun, F. (2024). In situ structural insights into the excitation-contraction coupling mechanism of skeletal muscle. Sci Adv 10, eadl1126. 10.1126/sciadv.adl1126.

7. Dixon, R.E., and Trimmer, J.S. (2022). Endoplasmic Reticulum-Plasma Membrane Junctions as Sites of Depolarization-Induced Ca(2+) Signaling in Excitable Cells. Annu Rev Physiol. 10.1146/annurev-physiol-032122-104610.

8. Bleck, C.K.E., Kim, Y., Willingham, T.B., and Glancy, B. (2018). Subcellular connectomic analyses of energy networks in striated muscle. Nat Commun 9, 5111. 10.1038/s41467-018-07676-y.

9. Kawaguchi, K., and Fujita, N. (2023). Shaping Transverse-Tubules: Central Mechanisms that Play a Role in the Cytosol Zoning for Muscle Contraction. J Biochem. 10.1093/jb/mvad083.

10. Smith, J.A.B., Murach, K.A., Dyar, K.A., and Zierath, J.R. (2023). Exercise metabolism and adaptation in skeletal muscle. Nat Rev Mol Cell Biol. 10.1038/s41580-023-00606-x.

11. Scorrano, L., De Matteis, M.A., Emr, S., Giordano, F., Hajnoczky, G., Kornmann, B., Lackner, L.L., Levine, T.P., Pellegrini, L., Reinisch, K., et al. (2019). Coming together to define membrane contact sites. Nat Commun 10, 1287. 10.1038/s41467-019-09253-3.

12. Jing, J., Liu, G., Huang, Y., and Zhou, Y. (2020). A molecular toolbox for interrogation of membrane contact sites. J Physiol 598, 1725–1739. 10.1113/JP277761.

13. Zhou, Z., Torres, M., Sha, H., Halbrook, C.J., Van den Bergh, F., Reinert, R.B., Yamada, T., Wang, S., Luo, Y., Hunter, A.H., et al. (2020). Endoplasmic reticulum-associated degradation regulates mitochondrial dynamics in brown adipocytes. Science 368, 54–60. 10.1126/science.aay2494.

14. Bryson-Richardson, R.J., and Currie, P.D. (2008). The genetics of vertebrate myogenesis. Nat Rev Genet 9, 632–646. 10.1038/nrg2369.

15. Gibbs, E.M., Horstick, E.J., and Dowling, J.J. (2013). Swimming into prominence: the zebrafish as a valuable tool for studying human myopathies and muscular dystrophies. FEBS J 280, 4187–4197. 10.1111/febs.12412.

16. Tesoriero, C., Greco, F., Cannone, E., Ghirotto, F., Facchinello, N., Schiavone, M., and Vettori, A. (2023). Modeling Human Muscular Dystrophies in Zebrafish: Mutant Lines, Transgenic Fluorescent Biosensors, and Phenotyping Assays. Int J Mol Sci 24. 10.3390/ijms24098314.

17. Peddie, C.J., Genoud, C., Kreshuk, A., Meechan, K., Micheva, K.D., Narayan, K., Pape, C., Parton, R.G., Schieber, N.L., Schwab, Y., et al. (2022). Volume electron microscopy. Nat Rev Methods Primers 2, 51. 10.1038/s43586-022-00131-9.

18. Micheva, K.D., Burden, J.J., and Schifferer, M. (2024). Array tomography: trails to discovery. Methods Microsc 1, 9–17. 10.1515/mim-2024-0001.

19. Takekura, H., Flucher, B.E., and Franzini-Armstrong, C. (2001). Sequential docking, molecular differentiation, and positioning of T-Tubule/SR junctions in developing mouse skeletal muscle. Dev Biol 239, 204–214. 10.1006/dbio.2001.0437.

20. Boncompagni, S., Rossi, A.E., Micaroni, M., Beznoussenko, G.V., Polishchuk, R.S., Dirksen, R.T., and Protasi, F. (2009). Mitochondria are linked to calcium stores in striated muscle by developmentally regulated tethering structures. Mol Biol Cell 20, 1058–1067. 10.1091/mbc.E08-07-0783.

21. Dirksen, R.T. (2009). Sarcoplasmic reticulum-mitochondrial through-space coupling in skeletal muscle. Appl Physiol Nutr Metab 34, 389–395. 10.1139/H09-044.

22. Eisner, V., Csordas, G., and Hajnoczky, G. (2013). Interactions between sarco-endoplasmic reticulum and mitochondria in cardiac and skeletal muscle - pivotal roles in Ca(2)(+) and reactive oxygen species signaling. J Cell Sci 126, 2965–2978. 10.1242/jcs.093609.

23. Du, S.J., Gao, J., and Anyangwe, V. (2003). Muscle-specific expression of myogenin in zebrafish embryos is controlled by multiple regulatory elements in the promoter. Comp Biochem Physiol B Biochem Mol Biol 134, 123–134. 10.1016/s1096-4959(02)00194-x.

24. Roostalu, U., and Strähle, U. (2012). In Vivo Imaging of Molecular Interactions at Damaged Sarcolemma. Developmental Cell 22, 515–529. 10.1016/j.devcel.2011.12.008.

25. Noble, S., Godoy, R., Affaticati, P., and Ekker, M. (2015). Transgenic Zebrafish Expressing mCherry in the Mitochondria of Dopaminergic Neurons. Zebrafish 12, 349–356. 10.1089/zeb.2015.1085.

26. Cali, T., and Brini, M. (2021). Quantification of organelle contact sites by split-GFP-based contact site sensors (SPLICS) in living cells. Nat Protoc 16, 5287–5308. 10.1038/s41596-021-00614-1.

27. Vallese, F., Catoni, C., Cieri, D., Barazzuol, L., Ramirez, O., Calore, V., Bonora, M., Giamogante, F., Pinton, P., Brini, M., and Cali, T. (2020). An expanded palette of improved SPLICS reporters detects multiple organelle contacts in vitro and in vivo. Nat Commun 11, 6069. 10.1038/s41467-020-19892-6.

28. Giamogante, F., Barazzuol, L., Poggio, E., Tromboni, M., Brini, M., and Cali, T. (2022). Stable Integration of Inducible SPLICS Reporters Enables Spatio-Temporal Analysis of Multiple Organelle Contact Sites upon Modulation of Cholesterol Traffic. Cells-Basel 11. 10.3390/cells11101643.

29. Lee, E., Marcucci, M., Daniell, L., Pypaert, M., Weisz, O.A., Ochoa, G.C., Farsad, K., Wenk, M.R., and De Camilli, P. (2002). Amphiphysin 2 (Bin1) and T-tubule biogenesis in muscle. Science 297, 1193–1196. 10.1126/science.1071362.

30. Smith, L.L., Gupta, V.A., and Beggs, A.H. (2014). Bridging integrator 1 (Bin1) deficiency in zebrafish results in centronuclear myopathy. Hum Mol Genet 23, 3566–3578. 10.1093/hmg/ddu067.

31. Snead, W.T., Zeno, W.F., Kago, G., Perkins, R.W., Richter, J.B., Zhao, C., Lafer, E.M., and Stachowiak, J.C. (2019). BAR scaffolds drive membrane fission by crowding disordered domains. Journal of Cell Biology 218, 664–682. 10.1083/jcb.201807119.

32. Liu, T., Stephan, T., Chen, P., Keller-Findeisen, J., Chen, J., Riedel, D., Yang, Z., Jakobs, S., and Chen, Z. (2022). Multi-color live-cell STED nanoscopy of mitochondria with a gentle inner membrane stain. Proc Natl Acad Sci U S A 119, e2215799119. 10.1073/pnas.2215799119.

33. Katti, P., Hall, A.S., Parry, H.A., Ajayi, P.T., Kim, Y., Willingham, T.B., Bleck, C.K.E., Wen, H., and Glancy, B. (2022). Mitochondrial network configuration influences sarcomere and myosin filament structure in striated muscles. Nat Commun 13, 6058. 10.1038/s41467-022-33678-y.

34. Membranes, C.T.i. (2013). T-Tubule.

35. Hong, T., and Shaw, R.M. (2017). Cardiac T-Tubule Microanatomy and Function. Physiol Rev 97, 227–252. 10.1152/physrev.00037.2015.

36. Nelson, S.R., Li, A., Beck-Previs, S., Kennedy, G.G., and Warshaw, D.M. (2020). Imaging ATP Consumption in Resting Skeletal Muscle: One Molecule at a Time. Biophys J 119, 1050–1055. 10.1016/j.bpj.2020.07.036.

37. Ackers-Johnson, M., Li, P.Y., Holmes, A.P., O’Brien, S.M., Pavlovic, D., and Foo, R.S. (2016). A Simplified, Langendorff-Free Method for Concomitant Isolation of Viable Cardiac Myocytes and Nonmyocytes From the Adult Mouse Heart. Circ Res 119, 909–920. 10.1161/circresaha.116.309202.

